# About samples, giving examples: Optimized Single Molecule Localization Microscopy

**DOI:** 10.1101/568295

**Authors:** Angélique Jimenez, Karoline Friedl, Christophe Leterrier

**Affiliations:** Aix Marseille Université, CNRS, INP UMR7051, NeuroCyto, Marseille, France; Abbelight, Paris, France

**Keywords:** super-resolution microscopy, SMLM, STORM, DNA-PAINT, cytoskeleton

## Abstract

Super-resolution microscopy has profoundly transformed how we study the architecture of cells, revealing unknown structures and refining our view of cellular assemblies. Among the various techniques, the resolution of Single Molecule Localization Microscopy (SMLM) can reach the size of macromolecular complexes and offer key insights on their nanoscale arrangement in situ. SMLM is thus a demanding technique and taking advantage of its full potential requires specifically optimized procedures. Here we describe how we perform the successive steps of an SMLM workflow, focusing on single-color Stochastic Optical Reconstruction Microscopy (STORM) as well as multicolor DNA Points Accumulation for imaging in Nanoscale Topography (DNA-PAINT) of fixed samples. We provide detailed procedures for careful sample fixation and immunostaining of typical cellular structures: cytoskeleton, clathrin-coated pits, and organelles. We then offer guidelines for optimal imaging and processing of SMLM data in order to optimize reconstruction quality and avoid the generation of artifacts. We hope that the tips and tricks we discovered over the years and detail here will be useful for researchers looking to make the best possible SMLM images, a pre-requisite for meaningful biological discovery.

## Introduction

Optical microscopy of immunofluorescence-labeled samples has revolutionized biology by allowing access to cellular processes in their native setting. However, the resolution of an optical microscope is physically limited to about 200 nanometers due to the diffraction of light that occurs along the optical path (Hecht, 2015; Vangindertael et al., 2018). This limit prevents the detailed visualization of key cellular structures: organelles (mitochondria, endosomes), cytoskeleton assemblies (actin, microtubules, intermediate filaments) and other scaffolding structures such as clathrin-coated pits that all have typical dimensions between 10 and 500 nm (Milo and Phillips, 2015). New optical techniques, collectively called super-resolution microscopy, can now overcome this limit and resolve details down to a few tens of nanometers (Schermelleh et al., 2019). Since its emergence in the beginning of the 2000s, super-resolution microscopy has matured and is now used in many laboratories to investigate the nanoscale cellular architecture (Sahl et al., 2017; Sigal et al., 2018).

Among the widely available super-resolutive techniques, Single Molecule Localization Microscopy (SMLM) is the one that can attain the finest precision, getting close to ultrastructural details of a few nanometers in the best cases (Sieben et al., 2018b). SMLM obtains information beyond the diffraction limit by pinpointing the position of single fluorophores, a mechanism that is distinct from other strategies such as STimulated Emission Depletion (STED) or Structured Illumination Microscopy (SIM) (Vangindertael et al., 2018). Experimental comparisons of super-resolution methods can be found in (Bachmann et al., 2016) and (Wegel et al., 2016). Due to diffraction by the microscope objective, a single fluorophore emitting light will appear as a ∼200 nm-wide spot called the point-spread function (PSF). However, if the fluorophore is well isolated, it is possible to fit its position with a precision well beyond the size of the PSF (Fig. 1A). This localization precision primarily depends on the number of photons emitted by the fluorophores and is typically around ∼8 nm for standard deviation (s.d., 20 nm in FWHM) (Mortensen et al., 2010; Thompson et al., 2002). Single-molecule localization was thus used since the 1980s to track the movement of single particles and applied to fluorescence imaging to follow the diffusion of membrane proteins (Schmidt et al., 1996) or the movement of molecular motors (Yildiz, 2003).

**Figure 1.**
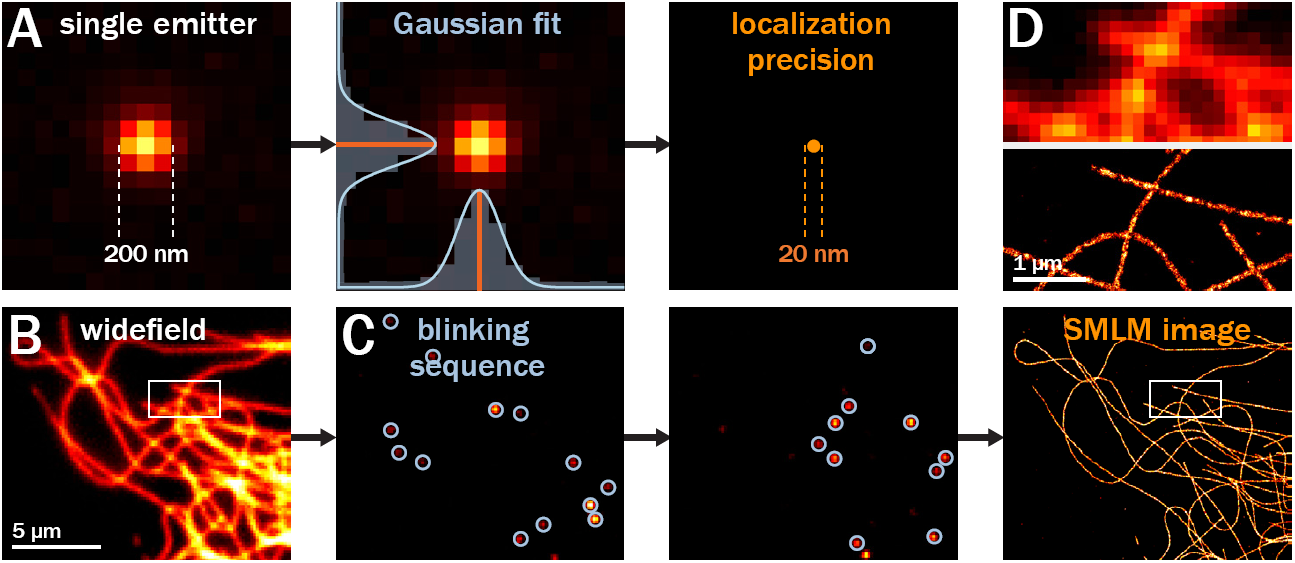
Principle of Single Molecule Localization Microscopy (SMLM) **A.** Epifluorescence image of a single emitter, showing the ∼200 nm width of the PSF (left) that is fitted using Gaussian curves (center) to determine its position with a ∼20 nm precision (right). **B.** Epifluorescence image of microtubules in a COS cell (top) and corresponding cartoon (bottom). **C.** During SMLM acquisition, a blinking mode of fluorescence meission is induced and thousands of frames are recorded, containing individual blinking events that can be fitted to localize each emitter. **D.** After processing, all localizations are plotted to generate the SMLM images (bottom). Top panel is a zoom corresponding to the box highlighted in the full image and shows the gain in resolution with much thinner microtubules (top).

To take advantage of single-molecule localization and generate an image of a fluorescently labeled sample, it is necessary to separate the PSF from each fluorophore, as initially proposed by Betzig (Betzig, 1995). SMLM separates fluorophores along the temporal dimension by photoswitching: each fluorophore labeling the sample is active only very briefly, emitting fluorescence as one or several short spontaneous blinking events. At any given time, most fluorophores are dark and only a few of them are blinking, allowing to localize them by fitting their position (Fig. 1B). The blinking of fluorophores is generally acquired as a long sequence of thousands of frames, in order to localize millions of fluorophores. The SMLM image is then reconstructed by plotting all the localized fluorophores, resulting in a super-resolved image that has a ∼10X better resolution than the diffraction-limited image (Fig. 1C).

The mechanism used to generate blinking fluorophores distinguishes several techniques that belong to SMLM: PALM, STORM and DNA-PAINT. In Photoactivated Localization Microscopy (PALM), photoactivatable or photoconvertible fluorescent proteins are used and blinking is generated by illuminating the sample to sparsely activate or convert these fluorescent proteins (Betzig et al., 2006; Hess et al., 2006). One advantage of PALM is that it can be performed on living cells expressing photoactivatable protein fusions. This notably allows to access the dynamics of single proteins, in a variant called sptPALM (Manley et al., 2008). In STochastic Optical Reconstruction Microscopy (STORM), high intensity illumination and an oxygen-deprived, reducing buffer is generally used to induce sparse blinking of organic fluorophores commonly used for immunolabeling (Heilemann et al., 2008; Rust et al., 2006). Single-color STORM is readily compatible with classical immunolabeled samples, by using Alexa Fluor 647-coupled secondary antibodies (Halpern et al., 2015). Multi-color STORM is not as straight-forward, as it is a challenge to find two distinct fluorophores that have good blinking characteristics in the same environment, and inducing photoswitching of fluorophores outside of the far-red channel usually require high laser power illumination (Dempsey et al., 2011; Lehmann et al., 2016). In DNA Points Accumulation for imaging in Nanoscale Topography (DNA-PAINT) (Molle et al., 2016; Sharonov and Hochstrasser, 2006), blinking is generated by the transient hybridization of short DNA single-strand coupled to a fluorophore (imager strand) with its complementary strand (docking strand) attached to an antibody targeting the structure of interest (Jungmann et al., 2014; Nieves et al., 2018). DNA-PAINT does not require a specific buffer or very high laser power, as the blinking does not depend on the photophysics of the fluorophore. Moreover, fluorophores are constantly renewed at the sample as new imagers interact with the docking strands, allowing to accumulate a large number of localizations. Finally, multi-color DNA-PAINT is straightforward by using orthogonal docking strands on distinct secondary antibodies and corresponding imagers, allowing to image 5-8 different targets (Agasti et al., 2017; Almada et al., 2019; Yu Wang et al., 2017; Werbin et al., 2017).

Since their invention, these different SMLM techniques have been extensively used to probe the nanoscale arrangement of cellular structures and macromolecular complexes (Baddeley and Bewersdorf, 2018; Liu et al., 2015). Nevertheless, SMLM is a challenging technique that requires specific knowledge and skills in order to obtain good quality data (Lambert and Waters, 2017). We have been using STORM since 2013, then DNA-PAINT to study the organization of the axonal cytoskeleton (Leterrier et al., 2017; Papandréou and Leterrier, 2018), resolving the architecture of axonal actin and spectrins (Ganguly et al., 2015; C. Y.-M. Huang et al., 2017; Leterrier et al., 2015) as well as the mechanisms of slow axonal transport (Chakrabarty et al., 2019; Ganguly et al., 2017). We have also helped developing SMLM imaging modalities (Almada et al., 2019) and analysis strategies (Culley et al., 2018; Laine et al., 2019). During these years, we have refined our SMLM workflow by optimizing sample preparation, imaging and analysis. In this Methods article, we aim at summarizing our experience to help researchers that use SMLM or are interested in trying it. We will provide advice and tips along the whole workflow of cell culture, fixation, immunolabeling, imaging and processing (Fig. 2). We will describe how to prepare good reference test samples for STORM and DNA-PAINT by labeling abundant targets in fibroblasts cells (Fig. 2A-B). Cytoskeleton (actin and microtubules) is a good SMLM benchmark target, as it forms well-known cellular patterns and can be labeled at high density. Other targets include clathrin-coated pits, which ∼100 nm size is ideal to verify and benchmark the proper performance of an SMLM setup. We will summarize the specific procedures and tweaks we use during imaging and briefly explain our processing workflow (Fig.2 C-D). This will complement the existing reviews and methodological articles on how to perform STORM and DNA-PAINT (Allen et al., 2013; de Linde et al., 2011; Halpern et al., 2015; Hoess et al., 2018; Pereira et al., 2015; Schnitzbauer et al., 2017; J. Xu et al., 2017). We hope this will be useful to the microscopy community and help interested researchers to get the best possible quality for their SMLM images.

**Figure 2.**
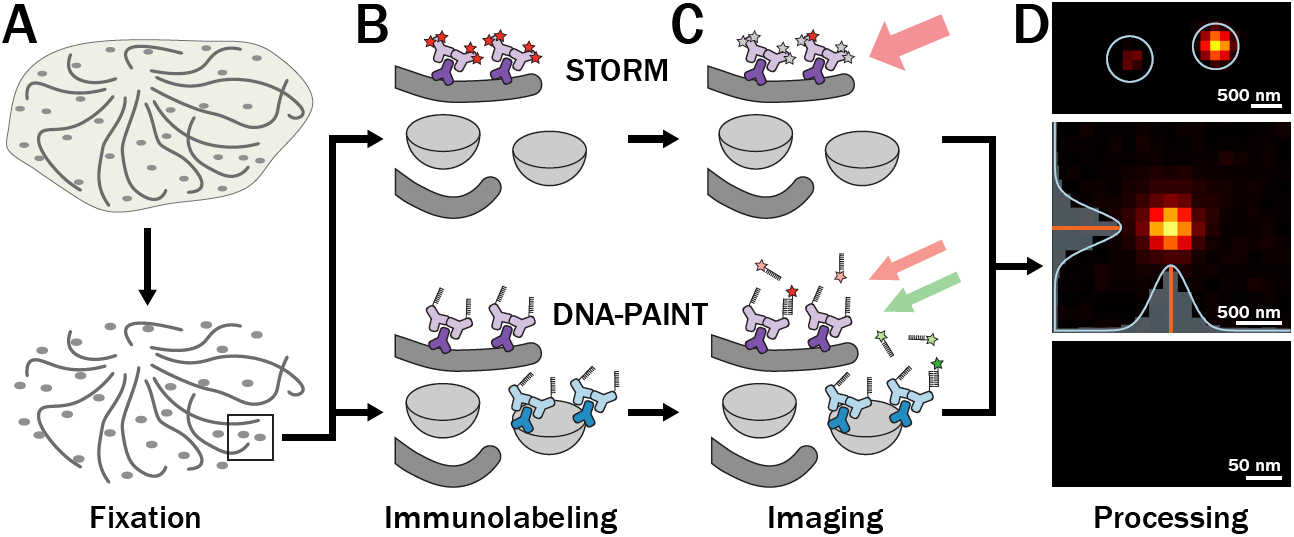
SMLM workflow. **A.** Cells are fixed and permeabilized. **B.** Cells are labeled using primary and secondary anti-bodies against structures such as microtubules (dark gray) and clathrin-coated pits (light gray). For STORM (top), secondary antibodies are conjugated with fluorophores. For DNA-PAINT (bottom), secondary antibodies are conjugated to short DNA single strands. **C.** STORM (top) uses high intensity illumination in a reducing buffer to induce fluorophore blinking. In DNA-PAINT (bottom), blinking occurs when imager strands in the medium interact with the docking strands. **D.** Processing includes detection of single emitters peaks (top), gaussian fitting of detected events (middle) and reconstruction using all localized events (bottom panel).

## Sample preparation

### Cell Culture

Flat and large cells make nice sample for SMLM, as they provide areas of thin cytoplasm rich in cytoskeletal and scaffolding elements. We are using COS-7 fibroblasts from Green African Monkey (ATCC, ref. CRL-1651) that are easy to culture and grow rapidly (Gluzman, 1981), but other mammalian cell lines are also typically used such as U2OS (Li et al., 2018), BS-C-1 (Bates et al., 2007; Lin et al., 2018), or HeLa, which are thicker and can be more challenging to image (Gao et al., 2018). It is important to know a bit about the underlying biology of the chosen cell line (use as a biological model, tumor origin…) in order to understand the possible singularities in their organization. For example, COS-7 cells frequently fail to properly divide and have a significant population of multi-nucleated cells. Sterile technique and health and safety regulations require a dedicated space for cell culture, and particular care must be taken to avoid contamination, notably by mycoplasmas, and by other cell lines leading to cell mis-identification (Geraghty et al., 2014). Proper cell culture technique should allow cells to grow over tens of passages without the need for constant antibiotic presence in the culture medium, and the culture should be regularly re-started from frozen stocks to avoid genetic drift and change in their properties.

### Cell seeding

The first step in SMLM sample preparation is to seed cells on glass coverslips. The best optical quality is obtained by using coverslips of 0.17 mm thickness, known as #1.5, for which high numerical aperture oil objectives (60X to 100X) typically used in SMLM are optimized. Some aspects of SMLM, notably 3D calibration, are highly dependent on coverslip thickness. In order to minimize variation from the calibrated curve, it is good to use high-precision thickness coverslips which have a thickness tolerance of 175 ± 5 um rather than 175 ± 15 um. We use 18-mm diameter round coverslips (Dutscher, ref. 900556) that are well adapted to our chambers and sample mounting options (see below). Coverslips should be thoroughly cleaned by successive baths: we clean our coverslips in racks using successive nitric acid, water and absolute ethanol baths followed by 2 hours at 180°C (Kaech and Banker, 2006). Once cleaned, coverslips are treated with poly-L-lysine to favor cell attachment (Sigma, ref. P2636); clean coverslips should become hydro-philic, with an easy spreading of the polylysine solution over the whole coverslip. After rinses, cells are seeded to 10-20% confluency and incubated overnight for attachment and spreading. A low confluency helps obtaining a significant proportion of single cells after 24 hours that will give the best results by SMLM.

### Calibration coverslips

When preparing coverslips for cell seeding, we usually prepare a couple of extra coverslips, up to the polylysine treatment, that we subsequently store in phosphate buffer. These are used to prepare sister calibration coverslips by just incubating them 5 minutes with 0.1 µm Tetraspeck beads (Thermo Fisher, ref. T7279) diluted 1:100-1:800 depending on the desired density of beads. Beads coverslips are notably useful for 3D-STORM calibration (see below), to verify the proper alignment of the laser and field homogeneity or calibrate chromatic aberration correction as they are fluorescent in the commonly used channels (488, 561 and 647 nm lasers).

## Immunostaining

### Fixation

Immunocytochemistry usually begins with the fixation of cells using chemical fixatives. An optimized fixation protocol is crucial to minimize artifacts that will be revealed at the nanoscale (Whelan and Bell, 2015). For example, cold methanol fixation is widely used before staining for microtubules (Bulinski, 1980), but in our experience results in poor preservation of microtubule organization at the nanoscale. As a rule of thumb, methods that have been validated at the ultrastructural level in electron microcopy sample preparation protocols are most likely to give the best results. As described previously (Whelan and Bell, 2015), we found that washing steps should be avoided before fixation. For the cytoskeleton, the best fixation uses glutaraldehyde in a cytoskeleton-preserving buffer. This can be preceded by a quick extraction step that will remove soluble proteins just before fixation, notably actin and tubulin monomers. Our pre-extraction/fixation protocol uses a moderate concentration of glutaraldehyde. It is optimal for microtubules and actin labeling but is also compatible with imaging of clathrin-coated pits (Fig. 3A, Fig. 4, Fig. 5, Fig. 6A and Fig. 7). After glutaraldehyde-based fixation, a quenching step using NaBH4 (Sigma, ref. 213462) as a reducing agent is necessary and works better than the NH4Cl/glycine incubation that is sometimes used.

**Figure 3.**
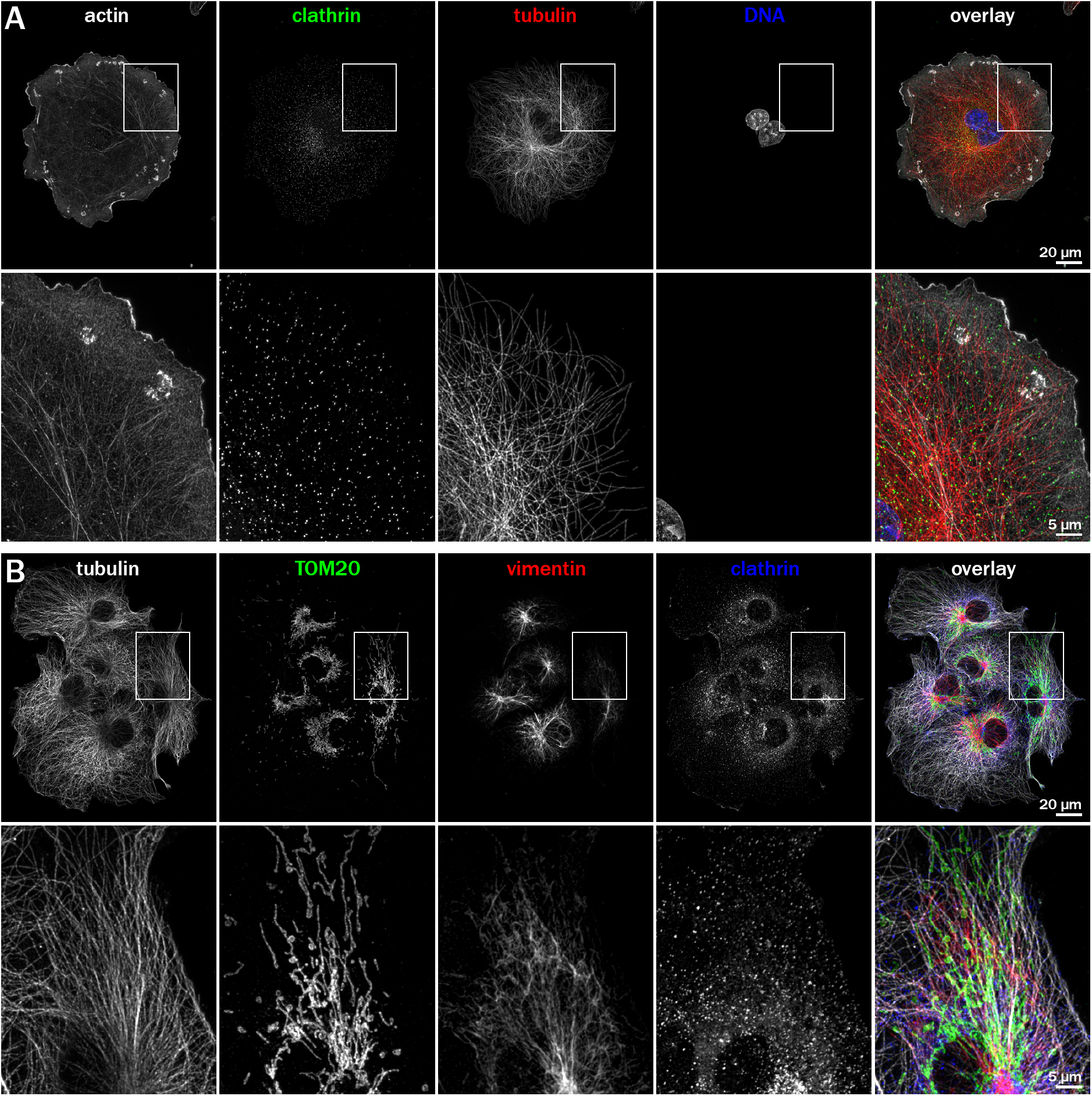
Diffraction-limited images of immunostained COS cells. Images are projections of deconvolved Z-stacks, acquired with an Apotome microscope (Zeiss) equipped with a 63X, NA 1.4 objective and a Flash4 v3 sCMOS camera (Hamama-tsu). **A.** COS cell fixed using the glutaraldehyde pre-extraction/fixation protocol, stained for actin (phalloidin, gray on overlay), microtubules (two anti-α-tubulin antibodies, green on overlay), clathrin (anti-clathrin heavy chain, red on overlay) and DNA (DAPI, blue on overlay). Bottom panels are zooms corresponding to the box highlighted on the full image (top panels). **B.** COS cells fixed using the hot paraformaldehyde fixation protocol, stained for microtubules (anti-tubulin, gray on overlay), intermediate filaments (anti-vimentin, red on overlay), mitochondria (anti-TOM020, green on overlay) and clathrin (anti-clathrin heavy chain, blue on overlay). Bottom panels are zooms corresponding to the box high-lighted on the full image (top panels).

**Figure 4.**
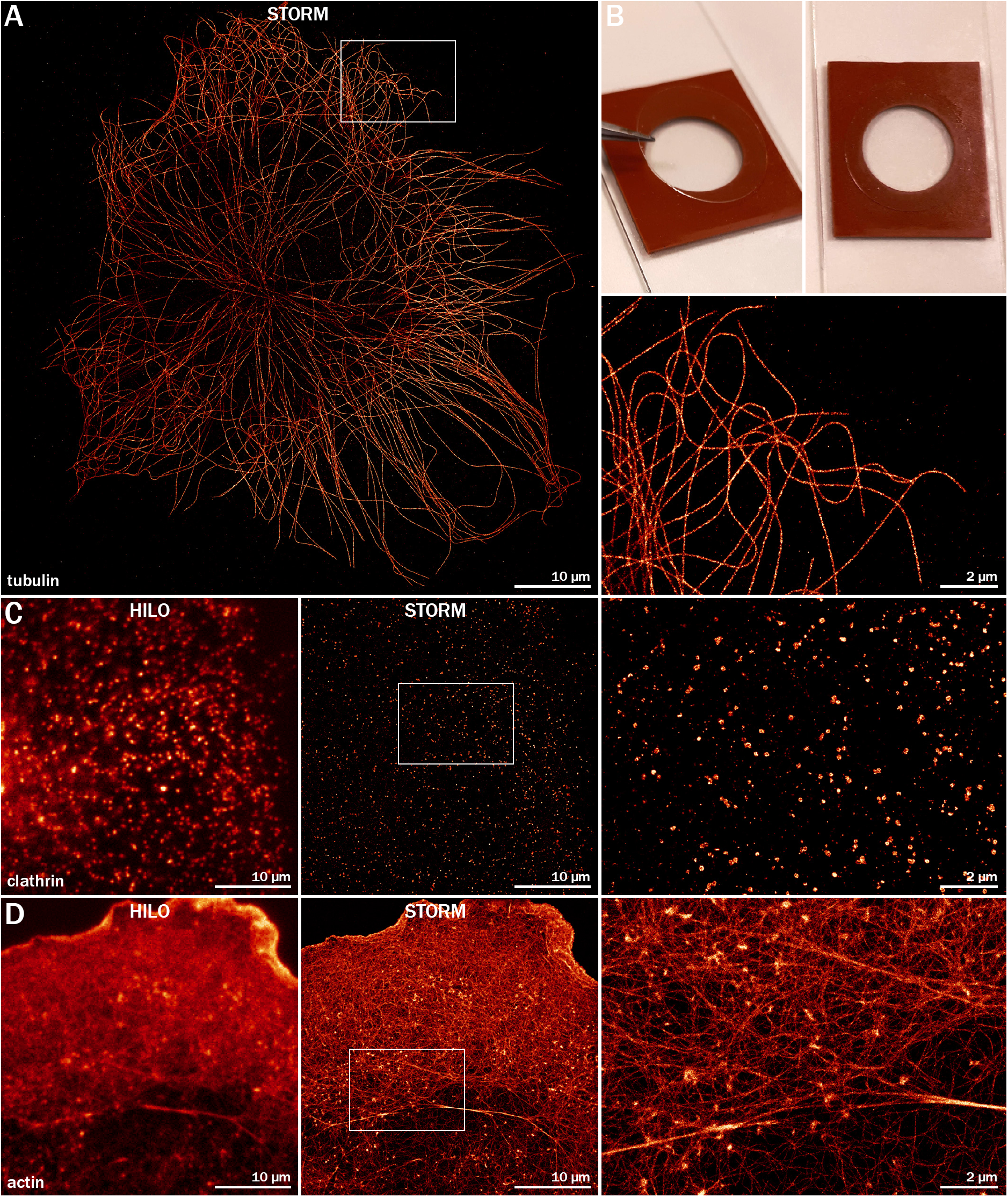
2D STORM images of microtubules, clathrin and actin in COS cells. **A**. STORM image of a COS cell labeled for microtubules (two anti-α-tubulin antibodies), acquired with an sCMOS camera (80 × 80 µm field of view). Right panel is a zoom of the area highlighted on the full image. **B**. Photograph showing the glass slide-attached silicone chamber (red) on which the 18 mm coverslip is sealed by gentle pression (left) to obtain a closed chamber filled with STORM buffer (right). **C**. Diffraction-limited HILO image (left panel) and corresponding STORM image (center panel) of a COS cell labeled for clathrin-coated pits (anti-clathrin light chain), acquired with an EMCCD camera (40 x 40 µm field of view). Right panel is a zoom of the area highlighted on the full image. **D**.Diffraction-limited HILO image (left panel) and corresponding STORM image (center panel) of a COS cell labeled for actin (phalloidin-AF647). Right panel is a zoom of the area highlighted on the full image.

**Figure 5.**
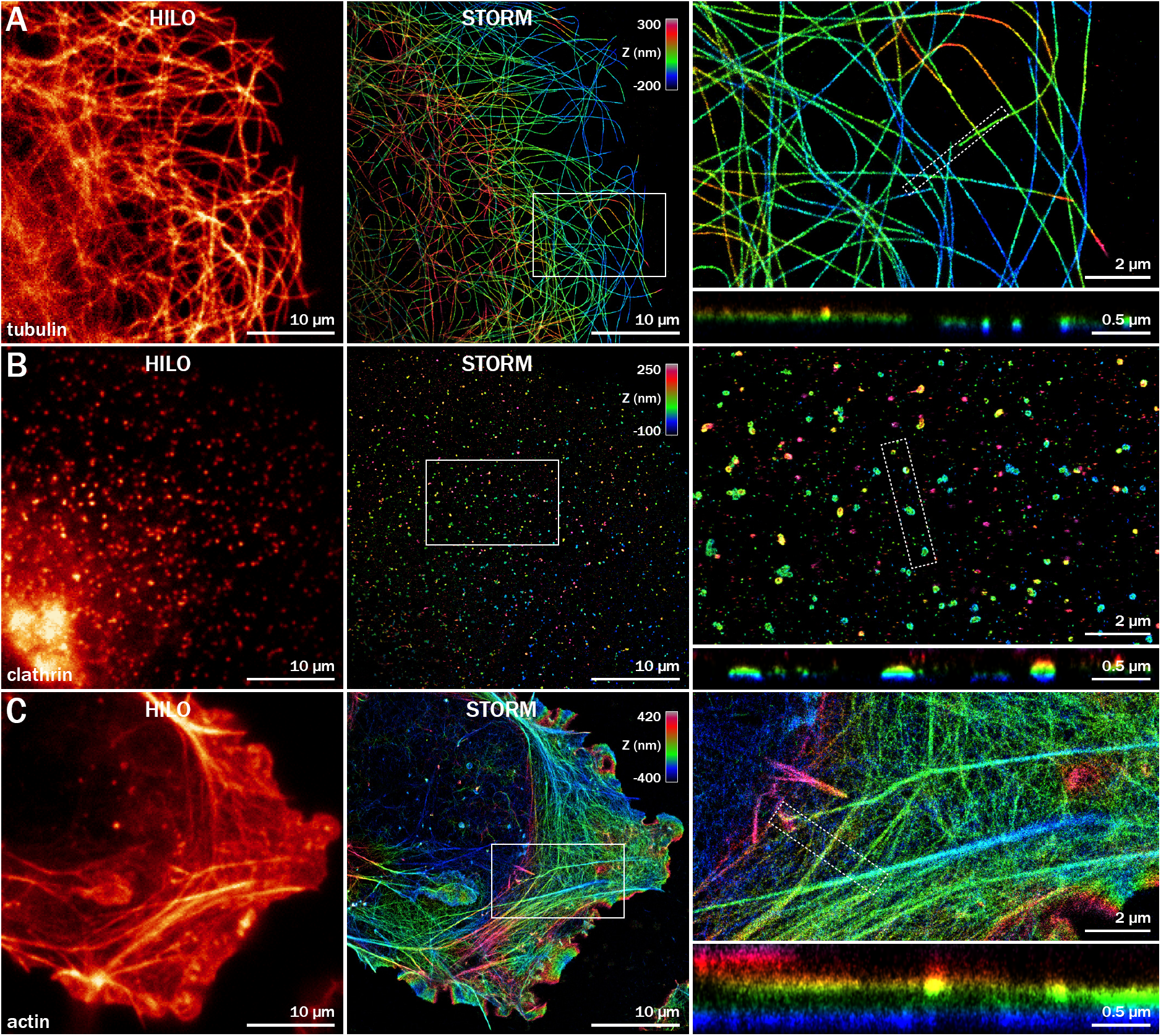
3D STORM images of microtubules, clathrin and actin in COS cells. **A.** Diffraction-limited HILO image (left panel) and corresponding astigmatism-based 3D STORM image (center panel) of a COS cell labeled for microtubules (two anti-α-tubulin antibodies), color-coded for depth. Top right panel is a zoom of the area highlighted on the full image, bottom right panel shows an XZ cross-section along the line highlighted on the zoomed image. **B.** Diffraction-limited HILO image (left panel) and corresponding 3D STORM image (center panel) of a COS cell labeled for clathrin-coated pits (anti-clathrin light chain), color-coded for depth. Top right panel is a zoom of the area highlighted on the full image, bottom right panel shows an XZ cross-section along the line highlighted on the zoomed image. **C.** Diffraction-limited HILO image (left panel) and corresponding 3D STORM image (center panel) of a COS cell labeled for actin (phalloidin-AF647), color-coded for depth. Top right panel is a zoom of the area highlighted on the full image, bottom right panel shows an XZ cross-section along the line highlighted on the zoomed image.

**Figure 6.**
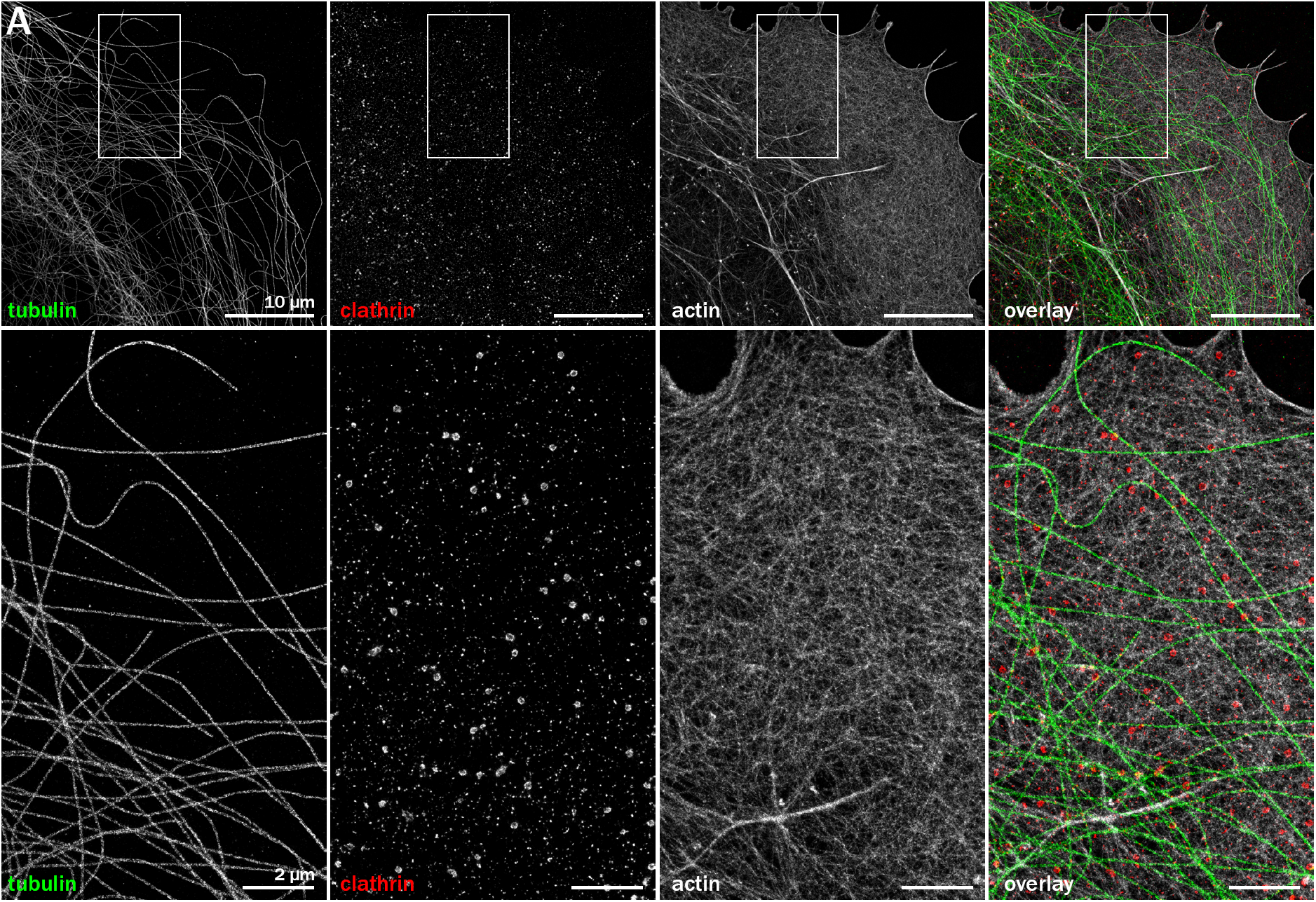
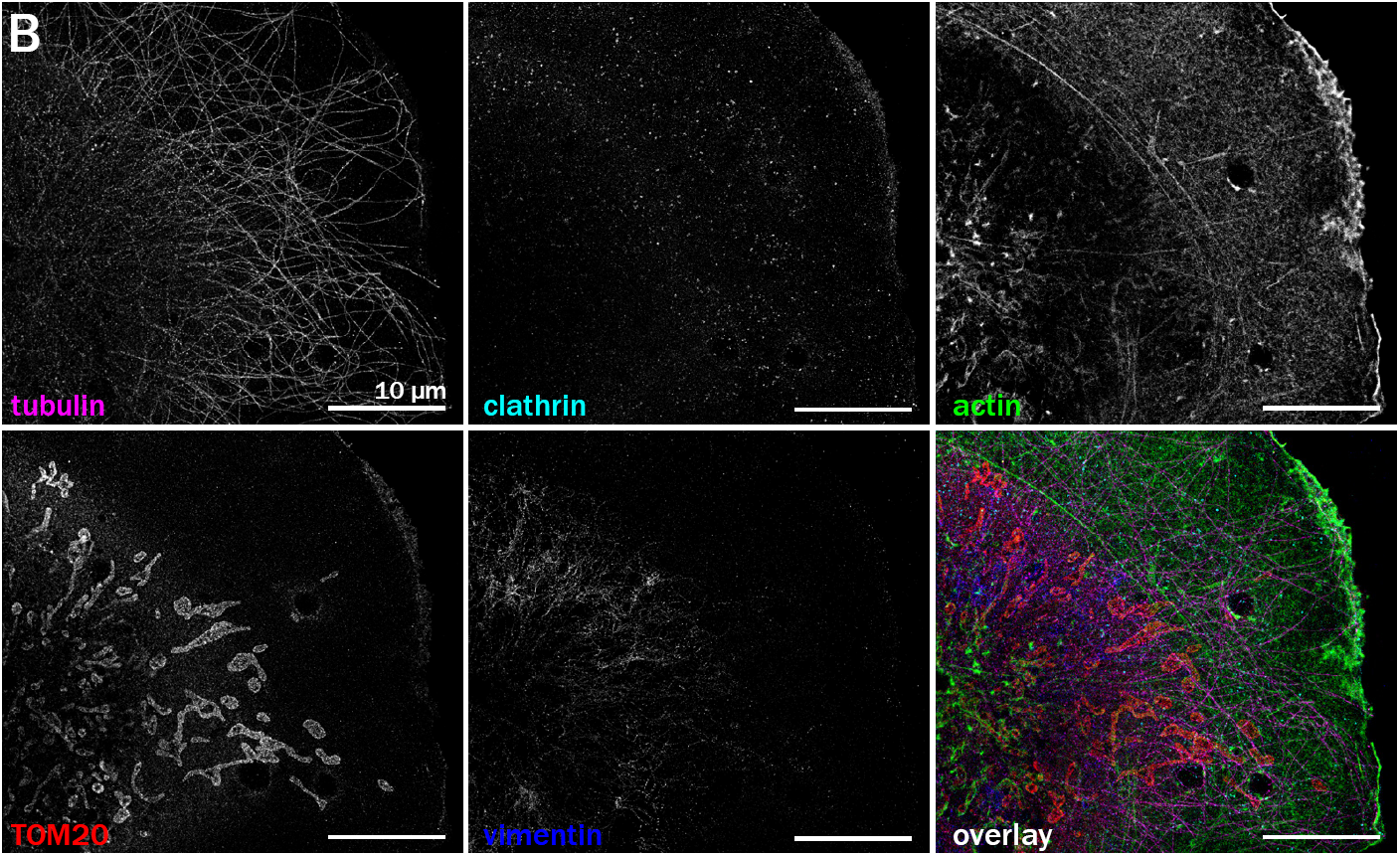
Multicolor STORM and DNA-PAINT images of immunostained COS cells. **A.** 3-color image obtained by sequential acquisition of actin by STORM (phalloidin-AF647, gray in overlay), microtubules (two anti-α-tubulin antibodies, green on overlay) and clath-rin-coated pits (anti-clathrin light chain, red on overlay) by DNA-PAINT (Atto655 imager). Bottom panels are zoomed images corresponding to the highlighted area on the full image. **B.** 5-color image obtained in three steps: first DNA-PAINT of actin (phalloidin-Atto488, green in overlay), microtubules (two anti-α-tubulin antibodies, Atto565 imager, magenta on over-lay) and mitochondria (anti TOM20, Atto655 imager, red on overlay), then DNA-PAINT of intermediate filaments (anti-vimentin, Cy3B imager, blue on overlay), then DNA-PAINT of clathrin (anti-clathrin light chain, Cy3B imager, cyan on overlay).

**Figure 7.**
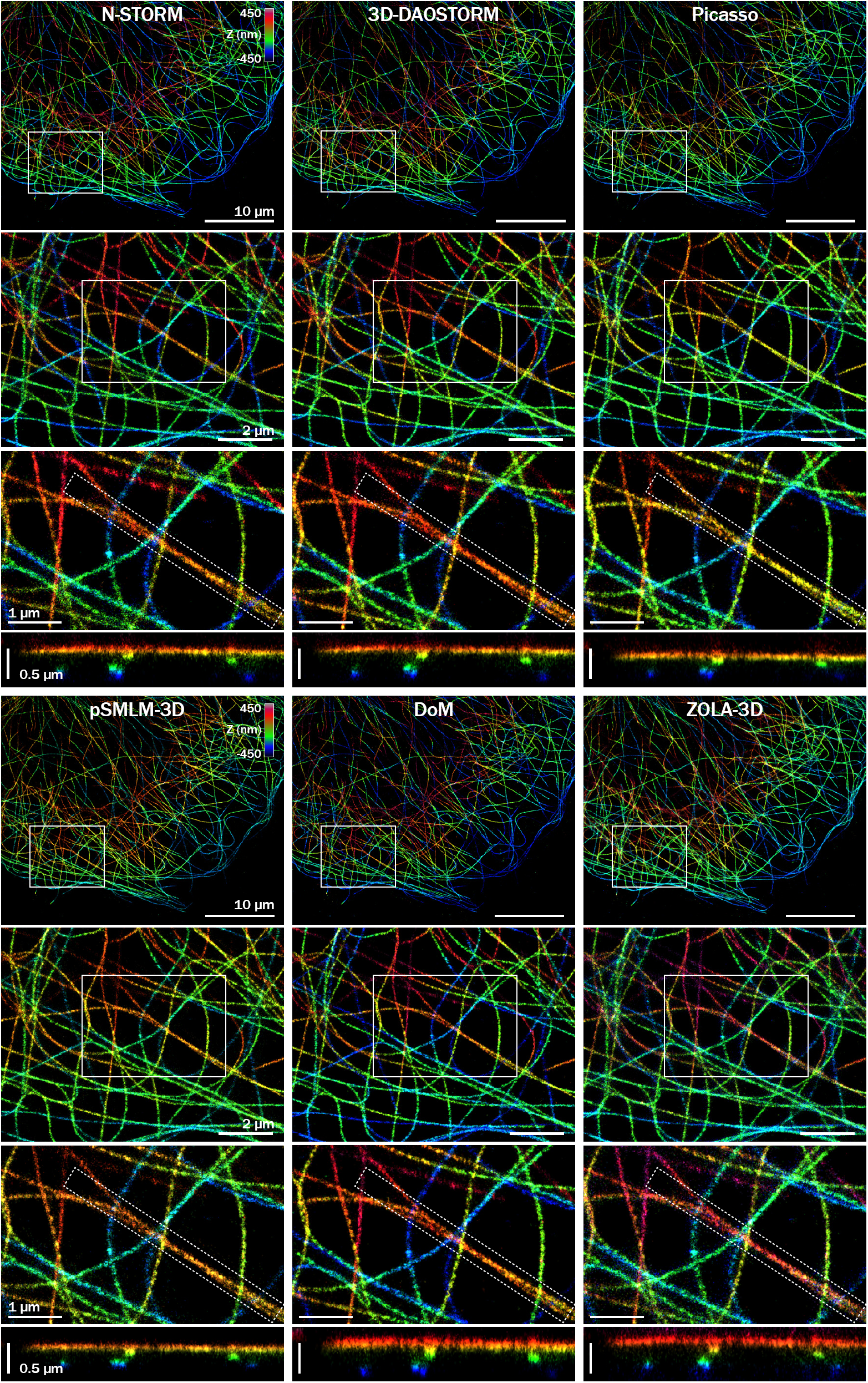
Processing of 3D-STORM data using several examples of SMLM software. COS cell labeled for microtubules (two anti-α-tubulin antibodies) and imaged by 3D STORM. The acquired image sequence has been processed using different software packages using optimized parameters for each software: N-STORM (Nikon, proprietary); 3D-DAOSTORM (open source, Python); Picasso (open source, Python); phasor pSMLM-3D (open source, ImageJ); DoM Utrecht (open source, ImageJ); ZOLA-3D (open source, ImageJ). See Table 2 for more information about each software.

#### Glutaraldehyde fixation protocol

PEM = 80 mM PIPES, 5 mM EGTA, 2 mM MgCl2, pH 6.8.

Extraction solution: 0.25% Triton, 0.1% glutaraldehyde in PEM, pre-heated to 37°C.

Fixation solution: 0.25% Triton, 0.5% glutaraldehyde in PEM, pre-heated to 37°C.

Work at 37°C using a water bath or pre-heated pad for the extraction and fixation steps.

- Incubate with extraction solution for 15 to 45 seconds.
- Replace with fixation solution and incubate for 10 minutes.
- Replace with 0.1% NaBH4 in phosphate buffer and incubate 7 minutes at room temperature.
- Rinse two times quickly with phosphate buffer.

**Table 1.** Primary antibodies used.

| Structure | Target protein | Species | Ig type | Clone | Supplier | Cat # | Dilution |
| --- | --- | --- | --- | --- | --- | --- | --- |
| microtubules | α-tubulin | rabbit polyclonal | – | – | abcam | ab18251 | 1:300 |
| microtubules | α-tubulin | mouse monoclonal | IgG1 | B-5-1-2 | Sigma | T 5168 | 1:300 |
| microtubules | α-tubulin | mouse monoclonal | IgG1 | DM1A | Sigma | T 6199 | 1:300 |
| microtubules | β-tubulin | mouse monoclonal | IgG1 | 2-28-33 | Sigma | T 5293 | 1:200 |
| clathrin-coated pits | clathrin heavy chain | rabbit polyclonal | – | – | abcam | ab21679 | 1:150 |
| microtubules | α-tubulin | rat monoclonal | – | YOL1/34 | abcam | ab6161 | 1:150 |
| microtubules | tyrosinated α-tubulin | rat monoclonal | – | YL1/2 | abcam | ab6160 | 1:150 |
| intermediate filaments | vimentin | chicken polyclonal | IgY | Poly 29191 | BioLegend | 919101 | 1:1000 |
| mitochondria | TOM20 | mouse monoclonal | IgG1 | 29/TOM20 | BD Bioscience | 612278 | 1:200 |

**Table 2.** Secondary antibodies used.

| Application | Host species | Target species | Fluorophore | Supplier | Cat # | Dilution | Concentration |
| --- | --- | --- | --- | --- | --- | --- | --- |
| STORM | donkey | mouse | Alexa Fluor 647 | Thermo Fisher | A31571 | 1:300 | 6.5 μg/mL |
| STORM | donkey | rabbit | Alexa Fluor 647 | Jackson ImmunoResearch | 711-605-152 | 1:300 | 5 μg/mL |
| DNA-PAINT | donkey | mouse | PAINT P1 sequence | Homemade |  | 1:100 |  |
| DNA-PAINT | donkey | rabbit | PAINT P3 sequence | Homemade |  | 1:100 |  |
| epifluorescence | donkey | rabbit | Dylight 405 | Rockland | 611-746-127 | 1:300 | 1.3 μg/mL |
| epifluorescence | donkey | mouse | Alexa Fluor 488 | Invitrogen | A21202 | 1:300 | 6.7 μg/mL |
| epifluorescence | goat | rat | Alexa Fluor 555 | Invitrogen | A21434 | 1:400 | 5 μg/mL |
| epifluorescence | donkey | rabbit | CF568 | Biotium | 20098 | 1:400 | 5 μg/mL |
| epifluorescence | goat | chicken | Alexa Fluor 647 | Invitrogen | A21449 | 1:300 | 6.7 μg/mL |

The glutaraldehyde-based fixation protocol will not allow optimal labeling for other targets such as mitochondria (TOM20/TOM22 protein) or intermediate filaments (vimentin). Alternatively, we use formaldehyde (FA) in cytoskeleton preserving buffer that works also very well for actin and clathrin (Leyton-Puig et al., 2016). If used at 37°C, this protocol can lead to satisfactory microtubule preservation (Fig. 3B, Fig. 7B). However, in our hands it is not robust and not as good as the glutaraldehyde-based fixation. Quenching after FA-based fixation is usually not necessary and does not provide a better immunolabeling. An important point for optimal sample preparation is the freshness of the aldehydes used for fixation: we use electron-microscopy grade glutaraldehyde (25%, Sigma, ref. G5882) and formaldehyde (37%, EMS Diasum, ref. 15714) in pure water, in glass ampoules. After opening, they are stored at 4°C and used within two weeks.

#### FA-PEM fixation

PEM = 80 mM PIPES, 5 mM EGTA, 2 mM MgCl2, pH 6.8.

Fixation solution: 4% FA, 4% sucrose in PEM.

- Fix for 10 minutes at room temperature or use fixation solution pre-heated to 37°C.
- Rinse three times quickly with phosphate buffer.

### Blocking and primary antibodies incubation

We found blocking to be an important step, and usually perform it for at least 1 hour at room temperature (and up to 3 hours when staining clathrin) with gentle agitation. We use gelatin as the blocking agent (Sigma, ref. G9391), and permeabilize the cells at the same time using 0.1% Triton X-100 (Sigma, ref. T8787). This blocking buffer (phosphate buffer 0.1 M pH 7.3, 0.22% gelatin, 0.1% Triton X-100) is also used for primary and secondary antibodies incubation. After blocking and permeabilization, we perform antibody incubations and rinses inside blackened-out large square Petri dishes, on Parafilm (Table 1). This allows to use a minimal amount of reagents: for antibodies and phalloidin incubations, 18 mm coverslips are flipped (cells facing downward) on 50 µL droplets and are flipped back (cells facing upward) for rinses.

For SMLM, it is important to maximize antibody coverage, i.e. ensure that the target epitopes are labeled with the highest possible efficiency. This entails using higher concentration of primary antibodies as long as this does not lead to overwhelming background (for example, anti-tubulin antibodies can be used at 10X the concentration used for regular epifluorescence labeling, Table 1). Moreover, higher labeling density for a given structure can be obtained by using several antibodies against different epitopes within this structure. This strategy has been used to maximize signal for synapses (Sigal et al., 2015) and microtubules in SMLM (Li et al., 2018) or Expansion Microscopy (Gao et al., 2018). Similarly, we usually label microtubules with a combination of anti-α- and anti-ß-tubulin monoclonal antibodies to obtain a dense labeling in STORM and PAINT (Fig. 3). Finally, in our hands the highest labeling density is obtained using overnight incubation of the primary antibody mixture at 4°C. In any case, it is always useful to optimize primary antibody incubation parameters (concentration, temperature, duration) depending on the type of sample and target.

### Secondary antibodies for STORM and PAINT

After primary antibodies incubation, the coverslips are rinsed using blocking buffer (3×10 minutes), then the cells are incubated with secondary antibodies in blocking buffer for 1 hour at room temperature. For STORM, we use commercial full IgG secondary antibodies coupled to Alexa Fluor 647 (Table 2) at a concentration of 4-7 µg/mL in blocking buffer. It is possible to co-stain cells using other antibodies revealed by different fluorophores such as Alexa Fluor 488 and Alexa Fluor 555. This allows to locate cells and interesting structures or complement the STORM image with diffraction-limited epifluorescence images from other targets. It is particularly useful for DNA-PAINT, because the DNA-coupled secondary antibodies do not provide steady-state fluorescence allowing to locate cells and features of interest. In this case, it is possible to label the same structure with an antibody against a distinct epitope and a fluorescent secondary antibody.

Labeling with primary and secondary antibodies (each with a ∼15 nm in size) adds a linkage error corresponding to the distance between the epitope and the fluorophore. In the case of microtubules, as antibodies can only go outward, this leads to a larger apparent diameter of ∼60 nm (versus 25 nm for the real external microtubule diameter). A solution to minimize linkage error is to use directly labeled primary antibodies or nanobodies (Mikhaylova et al., 2015). However, the use of direct labeling or nanobodies will result in less signal amplification and thus a reduced number of fluorophores and localizations (Pleiner et al., 2018), which can make the final structure appear spottier. In the general case where antibodies can bind on any side of the epitope, primary and secondary antibodies each add an uncertainty of s.d. ∼6.5 nm around the epitope position, which is then combined with the localization precision of the fluorophore. For typical STORM localization precision (s.d. ∼7 nm), this leads to a moderate degradation of the localization precision to s.d. ∼11 nm (FWHM from 16 to 25 nm) (Dani et al., 2010; Leterrier et al., 2015).

For multicolor imaging using the Exchange-PAINT variant of DNA-PAINT (Jungmann et al., 2014), we use specific secondary antibodies coupled to 9-basepair single strand of DNA (the “docking” strand). These DNA-coupled antibodies are not available commercially but can be prepared within two days in a biology lab thanks to published coupling protocols and sequences (Schnitzbauer et al., 2017). We are using the thiol coupling protocol as described in (Schnitzbauer et al., 2017) to conjugate donkey anti-mouse and donkey anti-rabbit antibodies with the P1 (ATACATCTA) and P3 (TCTTCATTA) docking strands, respectively. These secondary antibodies are tested for optimal concentration, which is usually 1:50-1:100 and incubated for 1 hour at room temperature in blocking buffer, similar to fluorescent secondary antibodies. Extra care should be taken to store the DNA-coupled antibodies, as the linked DNA strand tend to hydrolyze with time. We flash-freeze them in small aliquots just after coupling and keep the currently used aliquot at 4°C, using it within ∼1 month. After incubation, secondary antibodies are rinsed once with blocking buffer and twice with phosphate buffer (10 minutes each time).

### Actin staining

The best results for SMLM of actin are obtained using fluorescent phalloidin, because it allows for a very high density of labeling and results in crisp reconstruction of filamentous actin (K. Xu et al., 2012). As an alternative, PAINT of actin has been performed using phalloidin coupled to a DNA docking strand (Agasti et al., 2017) or actin-targeting affimers (Schlichthaerle et al., 2018a). It is also possible to use fluorescent LifeAct that transiently binds to actin in fixed samples to generate an SMLM image, a technique called Integrating exchangeable single-molecule localization (IRIS) (Ashdown et al., 2017; Kiuchi et al., 2015; Tas et al., 2018). In our hands, these alternatives are significantly slower than STORM with fluorescent phalloidin to generate a high-quality image.

For STORM, the best results are obtained using phalloidin-Alexa Fluor 647 (AF647, Thermo Fisher or Cell Signaling Technologies). An important point is that phalloidin-AF647 is more labile than uncoupled phalloidin or phalloidin coupled to other fluorophores, and tend to detach easily from actin, as noted previously (K. Xu et al., 2012). As a consequence, it is necessary to perform actin labeling as the very last step in the staining protocol. After secondary antibody rinses, cells are incubated in highly-concentrated phalloidin-AF647 (500 nM) in phosphate buffer, either for >1h at room temperature or overnight at 4°C. They are kept in phalloidin-AF647 until being imaged. When performing multi-color imaging with DNA-PAINT targets together with actin, we usually label actin with phalloidin-AF647 that gives the best results. Alternatively, we use phalloidin-Atto488 (500 nM phalloidin-Atto488 in phosphate buffer for >1h at room temperature) that can blink in PAINT imaging buffer (see below).

When followed precisely, this fixation and immunostaining protocol provides crisp and bright labeling of cellular structures. To verify the quality of sample preparation, we usually label sister coverslips with fluorescent secondary antibodies. After mounting on glass slides (Prolong Glass, Thermo Fisher), they can be imaged using regular epifluorescence or confocal microscopy to document the experiment (Fig. 3).

## Imaging

### Hardware setup

We perform SMLM using a Nikon N-STORM microscope equipped with a 100X, NA 1.49 TIRF objective, a Perfect Focus System for Z focus stabilization, an Andor 512×512 EMCCD camera and an Agilent laser launch (405 nm/25 mW, 488 nm/85 mW, 561 nm/85 mW, 647 nm/125 mW output). A 2X lens in the TIRF coupling arm focuses the laser to illuminate the center of the field of view, and SMLM acquisition is performed using the center 256×256 pixels (41 x 41 µm, 160 nm pixel size). A pixel size of 100 to 160 nm is generally used, corresponding to ≤1 s.d. of the PSF for reliable localization of single emitters (Thompson et al., 2002). The filter cube for STORM has four excitation bands corresponding to the 405, 488, 561 and 647 nm laser, and a single farred emission band (670-760 nm). When combining DNA-PAINT with STORM of actin labeled with phalloidin-Atto 488, we use a 4-band cube that has excitation and emission bands for each laser wavelength. Alternatively, we use a second setup consisting in an Abbelight module on an Olympus stand equipped with a 100X, NA 1.49 objective, Hama-matsu 2048×2048 sCMOS camera and an Oxxius laser launch (405 nm /100 mW, 488 nm/500 mW, 640 nm/500 mW). The larger sensor size allows for a smaller pixel size (97 nm) and a larger field of view (80 × 80 µm, Fig. 4A), requiring higher laser power to induce the fluorophore blinking necessary for STORM.

For optimal SNR, we use the laser illumination in the critical angle fluorescence microscopy mode (Nakata and Hirokawa, 2003; Sinkó et al., 2014), also called highly inclined and laminated optical sheet (HILO) (Tokunaga et al., 2008) or grazing incidence (Guo et al., 2018). This consists in the inclination of the laser angle until reaching the critical angle, a point where the sample is illuminated by a ∼1-2 µm thick horizontal light sheet above the coverslip, avoiding fluorescence from the upper part of the cell and imaging medium. This is perfectly adapted to the flat parts of the COS cells cytoplasm, and to the Z range of the astigmatism-based 3D SMLM (∼800 nm). In addition, intensity shows a ∼2X peak precisely at the critical angle (Carniglia et al., 1972) which further enhances signal and allows to precisely locate the critical angle using the live histogram of the acquired image.

### STORM

For STORM, we use a commercial glucose oxidase-based STORM buffer that allows for robust and reproducible performance (Abbelight Smart Kit dSTORM buffer). Alternatively, we prepare a home-made glucose-oxidase glucose oxidase/catalase/glucose oxygen capture system together with cysteamine (MEA) as a reducing agent:

#### STORM buffer

- 50 mM Tris pH8, 10 mM NaCl (from 2X stock stored at 4°C).
- 10% glucose (from 25% stock stored at 4°C, ThermoFisher, ref. 15023-021).
- 50 mM MEA (from 1 M stock in 0.36 M HCl Sigma, ref. 60-23-1). We prepare aliquots of solid MEA corresponding to ~1 mL of 1 M solution and store them at −20°C in a desiccator. 1 M solution is prepared extemporaneously and kept at 4°C for up to 1 week.
- 0.5 mg/mL glucose oxidase (Sigma, ref. G2133, from 20 mg/mL stock in 50/50 GOD buffer/glycerol stored at −20°C. GOD buffer is 24 mM PIPES, 4 mM MgCl2, 2 mM EGTA, pH 6.8).
- 40 µg/mL catalase (Sigma, ref. C40, from 5 mg/mL stock in 50/50 GOD buffer/glycerol stored at −20°C).

As this buffer will progressively acidify when in contact with ambient oxygen (Olivier et al., 2013; Swoboda et al., 2012), we use a closed chamber made with adhesive silicone inserts (CoverWell chambers, EMS Diasum #70334-A) attached to glass slide (Fig. 4B). The round 18 mm coverslip is simply pressed on the chamber overfilled with STORM buffer (∼160 µL), and suction between glass and silicone allows for good stability, even during long acquisitions. We have not monitored the pH evolution of our buffer during imaging, but blinking performance is usually stable for up to 4 hours in this sealed silicone chamber. Collapse of buffer performance is usually detected as a slowing down of blinking (longer blinking events) and an inability to drive enough fluorophores in the dark state (appearance of constant fluorescence). Coverslips can be easily mounted and unmounted from the silicone insert, allowing to reuse them over several imaging sessions, and we keep them in phosphate buffer, 0.02% sodium azide in sealed multi-well plates between sessions.

Once mounted, a suitable cell is located using epifluorescence, then the illumination is switched to the 647 nm laser at the critical angle incidence and maximum laser power. After a few seconds, most fluoro-phores are brought to a dark state and the live image shows fast blinking of the activated fluorophores. We then launch the acquisition of 40,000 to 100,000 frames with an exposure time of 15 ms (20,000 frames / 50 ms on the Abbelight setup), which results in a total acquisition time of 10 to 20 minutes. During acquisition, illumination with the 405 nm laser is started and progressively raised in order to compensate for the bleaching of fast-blinking fluorophores, and to promote the blinking of fluorophores previously brought to a long-lived dark state. A feedback procedure coupled to live processing of blinking localizations automatically adjust the 405 nm laser intensity to maintain a constant number of detected localizations per frame. For best results, the goal is to maximize the number of localizations obtained in the minimum amount of time. The limit for this is the density of blinking fluorophores that will allow proper localization by the fitting algorithm (Wolter et al., 2011). Depending on the intrinsic density of the labeled structure, the 405 nm activation can be adjusted to stabilize the blinking close to the optimal density during the acquisition. It is thus possible to obtain several millions of localizations within 15 minutes, delineating cellular structures such as microtubules and clathrin-coated pits with great detail (Fig. 4A, 4C).

Actin labeling using phalloidin is usually the most densely blinking sample, often too dense to properly fit single blinking events. In order to decrease the blinking intensity, we raise the MEA concentration to 100 mM in the STORM buffer. As mentioned above, stained coverslips are kept in 500 nM phalloidin-AF647 until imaging. After equilibration at room temperature and three quick rinses with TpO4, coverslips are mounted in STORM buffer for imaging. Moreover, we supplement the STORM buffer with a lower concentration of phalloidin-AF647 (50 nM) that mitigates the detachment of phalloidin-AF647 during the imaging session, and renews the fluorescent phalloidin on the sample (Fig. 4D) (Ganguly et al., 2015).

### 3D imaging

3D imaging is performed by inserting a cylindrical lens in the optical path in front of the camera. This results in the deformation of the point-spread function (PSF) into an ellipse of opposite orientation below and above the focal plane (B. Huang et al., 2008; Kao and Verkman, 1994). After calibration using beads deposited on a coverslip, the shape of the PSF ellipse can be used to fit the position of the blinking fluorophore in Z over a ∼800 nm range, with a precision 2-3X worse than the lateral localization precision, (s.d. ∼20 nm, FWHM 50 nm) (B. Huang et al., 2008). Due to the enlarged PSF and the added dimension, algorithm for 3D SMLM usually have optimal performance at densities below those attainable in 2D SMLM (Sage et al., 2019), so this has to be taken into account during acquisition. 3D STORM images can be represented as Z color-coded images or transverse XZ/YZ sections showing the 3D organization of structures such as microtubules, clathrin-coated pits and actin filaments (Fig. 5).

### DNA-PAINT

DNA-PAINT principle is entirely distinct from STORM, but the resulting acquired sequences are similar, with a series of images showing blinking fluorophores. Acquisition of DNA-PAINT images is simpler than STORM, thanks to the separation of the blinking mechanism from the photophysics of the probe and illumination parameters. For each channel to image, the fluorescent imager strand complementary to the docking strand (coupled to the desired secondary antibody) is added to a saline buffer (PBS with 500 mM NaCl, pH 7.2), and imaging is done with continuous illumination at the critical angle. Blinking events density can be easily adjusted by changing the concentration of the imager strand - we typically use 0.1 to 1 nM. Once set, this density is usually constant over the whole acquisition, although sometimes high-power illumination can photolyze binding sites during long acquisition, resulting in a decreasing blinking event density (Blumhardt et al., 2018).

One thing to keep in mind is that DNA-PAINT acquisition is slower than STORM. The transient interaction between the 9-basepair docking strand and imager strand can last up to several hundreds of milliseconds, so blinking is significantly slower than in STORM. Typical exposure times of PAINT images are in the range of 100-500 ms, with the accumulation of tens of thousands of frames taking up to a few hours for a single image (Jungmann et al., 2014; Schlichthaerle et al., 2018b). One way to alleviate this is to use FRET-PAINT (Auer et al., 2017; Lee et al., 2017) that allows to raise the imager concentration without obtaining an overwhelming background fluorescence. We are using a simpler method when imaging densely stained targets: raising the laser power to 60-70% in the PAINT imaging buffer results in faster bleaching of bound probes, shortening each blinking event and decreasing the overall blinking density. This allows for high density blinking at faster frame rates (30-50 ms exposure time). As the fluorophore bound to the imager strand is usually bleached before it detaches from the docking strand, this does not result in a significant loss of photons emitted or localization precision. We typically acquire sequences of 20,000 to 40,000 frames for PAINT images.

We use PAINT rather than STORM for multicolor imaging, as we found it difficult to get reliably good blinking using a second fluorophore spectrally distinct from Alexa Fluor 647 (Dempsey et al., 2011). For two-color PAINT, there are two possibilities on our setup. The first is to use two different fluorophores (Atto565 and Atto655) on each imager strand (complementary to the P1 and P3 docking strands), and to image in the presence of both imager strands. Excitation is alternated every frame between the 561 and 647 nm lasers using the 4-band cube, and the two channels are acquired in a single acquisition sequence. One advantage of this procedure is that drift occurs in parallel for both channels and that alignment is easier, although the chromatic aberration between the two spectrally-distinct channels has to be corrected (Erdelyi et al., 2013). The other possibility, that we tend to favor because of its lower crosstalk between channels, is to use the same fluorophore for each imager with sequential acquisition separated by washing out of the first imager strand. In this case, we use far-red imagers (Atto655) with the single-emission band cube that result in a better signal (Fig. 6A). Channels must be corrected independently for drift and subsequently aligned using common features in both channels, and the use of the same fluorophore alleviates the need for chromatic correction. Both procedures can be combined for more than two colors: using sequential acquisition with two passes of two imagers (distinct fluorophores), we are able to acquire 4-color images of different targets (Fig. 6B). For easier rinses and medium exchange during these complex sequential imaging protocols, we mount the 18 mm round coverslips in an open metal chamber (Ludin chamber, Life Imaging Services) for PAINT imaging, and have devised an automatized pump system in LEGO driven by an Arduino computer and open-source software (Almada et al., 2019).

Finally, PAINT can be sequentially combined with STORM, and this is useful to image actin in addition to antibody-labeled structures. If actin is labeled with phalloidin-AF647, it is imaged first in STORM buffer, before imaging the PAINT channels sequentially using far-red imagers. To avoid residual actin staining interfering with the PAINT images, a ∼10 minutes bleaching step of the phalloidin-AF647 in PAINT imaging buffer at maximum 647 nm laser power must be performed before adding the first far-red PAINT imager. This allows to obtain the best actin images and combine them with other targets such as microtubules and clathrin (Fig. 6A). Alternatively, we use phalloidin-Atto488 that is well separated from the far-red PAINT channel and can blink in regular phosphate buffer or PAINT imaging buffer because of spontaneous quenching of cyanine dyes by the tryptophan residue of phalloidin (Nanguneri et al., 2014). STORM of actin is acquired first, with no imager present, before the acquisition of the PAINT channels (Fig. 6B) (Almada et al., 2019). There is significant bleaching of the phalloidin-Atto488 over time during the STORM acquisition, and Atto488 is insensitive to 405 nm activation. This ultimately limits the final quality of the actin image compared to phalloidin-AF647.

## Processing

### Fitting of the blinking events into localization coordinates

As SMLM images come from mathematical fitting performed over acquired images, processing is an integral part of the imaging workflow. It is important to have a good grasp of how this processing is performed, in order to be able to optimize parameters and avoid artifacts. The primary and most important processing step is the fitting of the blinking events: each peak on images from the acquisition sequence is detected, then fitted to determine the precise localization of the event. Fitting can be done using different algorithms, the most common ones being gaussian fitting with maximum likelihood or least-square fitting (Deschout et al., 2014; Sage et al., 2015). 3D SMLM involves localization in Z by fitting the point-spread function shape to a calibration curve. For densely labeled samples, specific algorithms are available that are able to fit overlapping peaks (Holden et al., 2011; Mailfert et al., 2018; Sage et al., 2019). Detection threshold (to decide what will be fitted) and rejection parameters (to avoid fitting peaks that are not single molecule blinking events) are important and should be adjusted to the acquisition conditions.

SMLM localization algorithms are now quite mature, with several software showing good performance. Rigorous evaluations of performance on standardized benchmark tests have been published for a number of software options both for 2D and 3D-SMLM, providing an unbiased overview of the available options (Sage et al., 2019; 2015). We will mention a few that we have tested extensively and found to give qualitatively satisfactory results on our data. We use the Nikon N-STORM localization software that is based on 3D-DAOSTORM and is available as an open-source python software (Babcock et al., 2012). The Abbelight Neo software is also used to process images acquired with this setup. A strong open-source option is Picasso (Schnitzbauer et al., 2017), a Python software that integrates GPU-accelerated least-square fitting (Przybylski et al., 2017) with various post-processing options. Several options are available in the open-source ImageJ/Fiji ecosystem (de Linde, 2019): a popular choice is ThunderSTORM (Ovesny et al., 2014), in particular the newer version integrating the phasor fitting approach, which is very fast and exists as a Fiji update site (Martens et al., 2018). Another good option is DoM (Detection of Molecules) Utrecht, a fast plugin that integrates options such as chromatic aberration correction (Chazeau et al., 2016). A recently-developed alternative is ZOLA-3D, which can fit any PSF shape including the ones used for extended Z-range (Aristov et al., 2018). There are also MATLAB-based software solutions that have good performance, such as fit3dspline and SMAP (Li et al., 2018; Sage et al., 2019). When properly used with optimized parameters, these software can provide good images from 2D and 3D SMLM acquisitions (Fig. 7, Table 3).

**Table 3.**
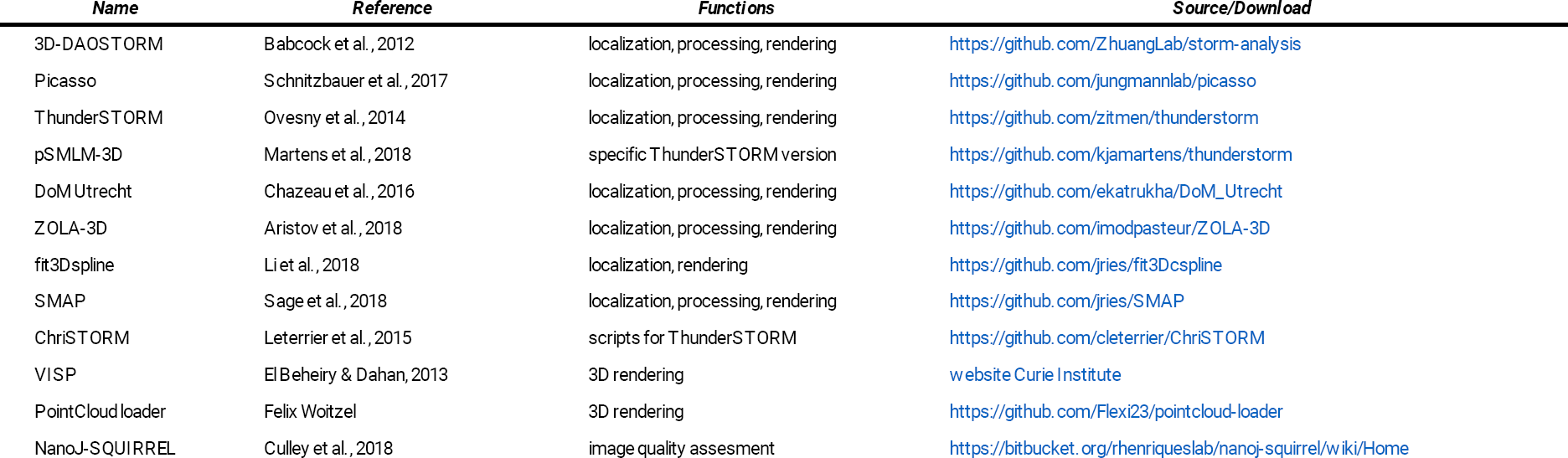
Software tools mentioned in the main text.

### Post-processing of localization coordinates

Once all blinking events have been fitted, several post-processing steps are necessary before reconstructing an image. The first one is drift correction: mechanical and thermal drift over tens to hundreds of nanometers occur during SMLM acquisitions, and proper drift correction is necessary to obtain a good image. For strongly structured images such as the cell cytoskeleton, the method of choice is the cross-correlation of images from parts of the acquisition sequence. Partial images are reconstructed using sub-sequences of ∼1000 frames and the drift is determined by cross-correlation with the preceding sub-sequence reconstruction, then interpolated within each frame of the sub-sequence. The best performance is obtained using redundant cross-correlation, where each sub-sequence reconstruction is correlated with all others (Yina Wang et al., 2014). It also possible to add fiducials to the sample, such as fluorescent or gold beads and use them to correct drift at the processing stage (Betzig et al., 2006), or to monitor drift during acquisition using brightfield imaging (McGorty et al., 2013). Drift correction is usually performed in 2D as a focus stabilization is used to avoid Z drift. However, Z drift can nevertheless occur (Cabriel et al., 2019) and can be further corrected by post-processing correction (3D-DAOSTORM, ZOLA-3D). Finally, sequential application of several drift correction steps can lead to progressively better results.

Another important post-processing step is to group together localizations from the same blinking event. Blinking events can sometimes last for two or more frames, and this is particularly true for DNA-PAINT acquisition where interaction between DNA strands can be long-lasting. In order to obtain a more faithful reconstruction with more precise localizations, this processing step groups together localizations that appear on subsequent frames within a given radius and averages the resulting localization (with a precision provided by photons from each frame). We usually set the radius to one camera pixel (160 nm on the N-STORM setup, 97 nm on the Abbelight setup). It is sometimes possible to specify a number of “dark” frames allowing transient disappearance of the blinking event.

Finally, it is important to filter the resulting localization list to remove localizations that are not good enough: one can filter on the least square fitting χ ^2^ (to exclude bad fits), the number of successive detections for grouped localizations (to avoid long-lasting events that are often artifacts), the number of photons emitted or the localization precision (to exclude blinking events that are too faint). Filtering on the size of the blinking event width can provide virtual optical sectioning by restricting the localizations to those very close to the focal plane (Palayret et al., 2015). Filtering the few localization events that have a bad precision can also speed up the reconstruction when using Gaussian rendering with individual precision values, as these localizations will contribute a low amount of intensity to a large number of pixels in the final image.

In our workflow, drift correction and localization grouping are done in the N-STORM software. When performing simultaneous two-color DNA-PAINT using red and far-red imagers, correction of the chromatic aberration is performed in the N-STORM software using a polynomial warp fitting calibrated on dense fields of Tetraspeck beads (Bates et al., 2012). After translation of the localization coordinates list in the proper comma-separated value (CSV) format, we do further post-processing in ThunderSTORM using custom scripts (ChriSTORM) (Leterrier et al., 2015), including another drift correction step and filtering of localizations. For STORM, we usually filter out localization events with less than 800 or more than 50,000 photons, and those lasting more than 5 successive frames. For PAINT, the upper limit is raised to 200,000 for photons emitted and to 50 for the number of successive frames.

### Image reconstruction

After fitting and post-processing, the final list of localization coordinates is the primary data format of SMLM. These can be used to reconstruct images, and several methods are available for this (Baddeley et al., 2010): histogram (the image is divided in a pixel grid and each pixel intensity depends on the number of localizations it contains), Gaussian rendering (a Gaussian is drawn at each localization coordinates), Delaunay triangulation (the image is tiled with one localization in each tile, the intensity being inversely proportional to the size of the tile). In the Gaussian case, the size of the Gaussian can be fixed (ideally to the average uncertainty of all localizations) or each can be drawn with its individual localization uncertainty depending on the number of photons emitted by the corresponding blinking event. The Gaussian rendering with individual precision values is more complex to generate but leads to the most precise reconstructions (Nieuwenhuizen and Stallinga, 2014). It is thus important for the fitting software to calculate, or at least approximate, the localization precision for each blinking event in order to use it for reconstruction. Finally, 3D STORM images can be reconstructed and visualized either as color-coded projections, or directly in 3D thanks to specific software such as ViSP (Beheiry and Dahan, 2013) or the web-based Pointcloud Loader (Table 3).

### Analysis of SMLM data

SMLM can provide fascinating image with extraordinary details of cellular structures. However, the primary data can (and should) be exploited beyond the reconstruction of images, preferably using coor-dinate-based analysis that directly use the localization list rather than the reconstructed image (Nicovich et al., 2017). This includes clustering and segmentation into structures, quantification of the nanoscale morphology, or relationships between distinct structures (Coltharp et al., 2014; Levet et al., 2015; Malkusch and Heilemann, 2016). At this scale, colocalization does not exist and is replaced by the measurement of distances between the localizations of different species (Georgieva et al., 2016; Lagache et al., 2018; Pageon et al., 2016). A recent development in SMLM analysis is the use of single particle averaging to reach molecular details beyond image resolution by averaging multiple similar objects (Broeken et al., 2015; Heydarian et al., 2018; Laine et al., 2015; Salas et al., 2017; Sieben et al., 2018a). A detailed review of the available algorithms and software is beyond the scope of the present article, but we encourage the interested reader to explore this very dynamic area that constitutes the next logical step toward leveraging SMLM possibilities. As an example, recent tools have been developed to assess the resolution and quality of SMLM images, allowing users to rapidly detect problems and optimize their imaging. Resolution in SMLM is tricky to define, as it involves not only the localization precision, but also the labeling density necessary to properly sample the structure of interest. Localization precision can be estimated over a single acquisition sequence by using localizations appearing on successive frames (Endesfelder et al., 2014), and Fourier ring correlation (FRC), an approach derived from single-particle electron microscopy, can provide resolution values that takes both aspects into account (Banterle et al., 2013; Nieuwenhuizen et al., 2013). Beyond resolution numbers, image quality can be assessed to detect artifacts and optimize any step of the SMLM workflow thanks to NanoJ-SQUIRREL, based on the comparison of the super-resolved image and its diffraction-limited counterpart (Culley et al., 2018).

## Conclusion

The exquisite resolution and access to single molecule measurements makes SMLM an attractive approach among the various super-resolution methods. It has brought crucial insights into the nanoscale organization of cells (Fornasiero and Opazo, 2015; Sigal et al., 2018). Commercial SMLM systems have been available for ten years, and they can be found in a lot of imaging facilities. However, SMLM is a technique that is still often considered as “difficult”: indeed, it requires careful planning and realization of the experiments, and each image takes time to acquire, in particular when performing multi-color PAINT experiments. In our experience, sample preparation is the most crucial point for good SMLM results and optimizing sample preservation while optimizing labeling density usually requires time and efforts. Processing principles must also be understood, in order to avoid the fabrication of artifacts (Burgert et al., 2015). Finally, taking advantage of single molecule data to obtain meaningful morphological and molecular insight is a current challenge, with a very active community developing new algorithms and software (Nicovich et al., 2017). We hope that our detailed workflow can serve as a good starting point and our advice help as pointers for further specific optimization.

## Author contributions, funding, and competing interests

AJ: acquisition of data, analysis and interpretation of data, revising the article. KF: acquisition of data, analysis and interpretation of data, revis-ing the article. CL: conception and design, acquisition of data, analysis and interpretation of data, drafting and revising the article. This work is supported by a grant to CL from CNRS (ATIP AO2016). KF is an employee of Abbelight.

## Acknowledgements

We thank Ghislaine Caillol, Fanny Boroni-Rueda, Sofia Yousfi and Martin Babeau for their help in preparing samples. We thank Pedro Pereira and Ricardo Henriques for DNA-PAINT secondary antibodies and help in preparing them.

